# CqHKT1 and CqSOS1 mediate genotype-dependent Na^+^ exclusion under high salt stress in quinoa

**DOI:** 10.1101/2024.08.05.606677

**Authors:** Yasufumi Kobayashi, Ryohei Sugita, Miki Fujita, Yasuo Yasui, Yoshinori Murata, Takuya Ogata, Yukari Nagatoshi, Yasunari Fujita

**Author notes:** Yasunari Fujita, Food Program, JIRCAS; 1-1 Ohwashi, Tsukuba, Ibaraki 305-8686, Japan, Corresponding author.

## Abstract

Salinity threatens crop production worldwide, and salinized areas are steadily increasing. As most crops are sensitive to salt, there is a need to improve the salt tolerance of major crops and promote the cultivation of under-utilized salt-tolerant crops. Quinoa, a pseudocereal and leafy vegetable from the Andean region of South America, is highly salt-tolerant, thrives in marginal environments, and has excellent nutritional properties. Research has often focused on epidermal bladder cells, a feature of quinoa thought to contribute to salt tolerance; however recent evidence suggests that these cells are not directly involved. The salt tolerance mechanism in quinoa remains unclear. Here, we show genotype-dependent differences in Na^+^ and K^+^ accumulation mechanisms using representative 18 lines of three genotypes by focusing on young quinoa seedlings at a stage without epidermal bladder cells. High salinity (600 mM NaCl) did not affect the early growth of all three quinoa genotypes. Under high salinity, lowland quinoa lines accumulated the most Na^+^ in the aerial parts, whereas southern highland lines accumulated the least. By contrast, K^+^ accumulation was slightly reduced in the aerial parts but significantly decreased in roots of all the genotypes. Resequencing of 18 quinoa lines supports the notion that genotype determines aboveground Na^+^ uptake and gene expression in response to salt stress. Using virus-induced gene silencing, we further demonstrated that CqHKT1 and CqSOS1 mediate Na^+^ exclusion in quinoa. These findings provide insight into salt tolerance mechanisms, serving as a basis for improving crop production under salt stress.

## Introduction

Salinity limits crop production in many regions of the world, and the affected areas and resulting economic losses are expected to increase (Melino and Tester 2023). Recently, not only has the FAO’s Global Map of Salt Affected Soils (GSASmap) covering 118 countries been released (FAO, 2021), but it has also been reported that there are 17 million square kilometers of salt-affected soils on the planet (Negacz et al. 2022). However, many crops, including staples such as maize (*Zea mays*), wheat (*Triticum aestivum*), rice (*Oryza sativa*), and soybean (*Glycine max*) are non-salt-tolerant plants (glycophytes). Salt-tolerant plants (halophytes) are extremely rare, comprising less than 0.25% of flowering plants (Bromham 2015). Therefore, there is a growing need to confer salt tolerance to conventional crops and to take advantage of salt-tolerant crops such as quinoa (*Chenopodium quinoa* Willd.). Deciphering the salt tolerance mechanisms of salt-tolerant plants may provide insights for improving salt tolerance in non-salt-tolerant crops (Melino and Tester 2023).

Quinoa, a facultative halophyte, is an annual C3 pseudocereal and leafy vegetable of the Amaranthaceae family, which also includes sugar beet (*Beta vulgaris* L.) and spinach (*Spinacia oleracea* L.). Quinoa is emerging as a potentially critical crop for global food and nutrition security due to its excellent nutritional value (Navruz-Varli and Sanlier 2016; Kobayashi et al. 2024) and ability to thrive in marginal environments (Bonifacio 2019). Cultivated in the Andes for over 7,000 years (Dillehay et al. 2007), quinoa was an essential part of the pre-Columbian Andean diet (González et al. 2015; Miller et al. 2021). After the Spanish conquest, quinoa cultivation was banned for religious reasons, in contrast to the global spread of other Andean crops such as tomatoes (*Solanum lycopersicum* L.) and potatoes (*Solanum tuberosum* L.), but it later regained interest in the 20th century (Gomez-Pando 2015; Ogata et al. 2021). The National Academy of Sciences (NAS) and the National Aeronautics and Space Administration (NASA) have recognized quinoa for its potential importance as a future food on and off the planet (NAS 1975; Schlick and Bubenheim 1993), and the United Nations declared 2013 the “International Year of Quinoa” (Bazile et al. 2016). Bolivia and Peru are the primary producers of quinoa, but this crop is now grown in over 120 countries (Alandia et al. 2020). Quinoa grows at a wide range of altitudes, from coastal areas to around 4,000 meters above sea level (Jacobsen 2003; Zurita-Silva et al. 2014; Bonifacio 2019). Quinoa can tolerate drought, high salinity, and frost, as exemplified by its cultivation near the Salar de Uyuni in Bolivia (Jacobsen 2003; Hariadi et al. 2011; Zurita-Silva et al. 2014; Yasui et al. 2016; Bonifacio 2019; Mizuno et al. 2020). However, growing quinoa outside the Andes poses challenges, due to its low tolerance of high temperatures, susceptibility to pests, and sensitivity to variations in day length (Gomez-Pando et al. 2019).

Plants have evolved not only morphological plasticity but also various physiological mechanisms, such as ion accumulation and exclusion, osmotic regulation, enhanced antioxidant responses, and ion homeostasis, that allow them to maintain growth under high salinity conditions (Munns and Tester 2008; Adolf et al. 2013; Deinlein et al. 2014; van Zelm et al. 2020). In terms of morphological characteristics, extensive research has explored how epidermal bladder cells on the leaf surface of salt-tolerant plants such as quinoa and common ice plants (*Mesembryanthemum crystallinum* L.), both of which are in the Caryophyllales order, contribute to salt stress tolerance (Agarie et al. 2007; Flowers and Colmer 2008; Yuan et al. 2016; Kiani-Pouya et al. 2017). Recent studies have shown that the salt tolerance of quinoa mutants lacking epidermal bladder cells is similar to that of the wild type and that K^+^ accumulates preferentially in quinoa epidermal bladder cells over Na^+^, indicating that epidermal bladder cells are directly involved in the salt tolerance mechanism (Moog et al. 2022). More recently, epidermal bladder cells in quinoa were implicated in herbivore defense mechanisms (Moog et al. 2023). These findings compel us to look beyond morphological features such as epidermal bladder cells in efforts to decipher the salt tolerance mechanism in quinoa. While abscisic acid (ABA)–mediated osmotic regulation (Fujita et al. 2011) and enhancement of antioxidant responses (Choudhury et al. 2017) are common responses to abiotic stresses such as drought and salinity (Zhu 2002; Golldack et al. 2014; Gupta et al. 2020), the mechanisms specifically required to respond for the plant’s response to salt stress are ion transport and ion homeostasis (Munns and Tester 2008; Deinlein et al. 2014; van Zelm et al. 2020). Molecular genetic and plant electrophysiological studies have shown that the ability to maintain high cytosolic K^+^/Na^+^ ratios is important for salt tolerance in plants to maintain ion homeostasis (Shabala and Cuin 2008). Transporters such as HKT1 and SOS1 are required for Na^+^ efflux from plants (Munns and Tester 2008; Deinlein et al. 2014; van Zelm et al. 2020). In the model plant Arabidopsis (*Arabidopsis thaliana*), the plasma membrane Na^+^/H^+^ antiporter SOS1 mediates root Na^+^ efflux and xylem Na^+^ loading, and the channel-like protein HKT1;1 functions in xylem Na^+^ unloading to regulate net Na^+^ uptake (Deinlein et al. 2014; van Zelm et al. 2020). A recent study showed that the Ca^2+^ sensor protein SOS3 inversely regulates the two major Na^+^ transporters SOS1 and HKT1 to mediate Na^+^ loading and unloading at the xylem in vascular plants, respectively (Gamez-Arjona et al. 2024). In quinoa, it remains to be determined whether transporters such as SOS1 and HKT1 play a role in salt tolerance, as observed in model plant studies.

Quinoa is an allotetraploid with 2*n* = 4*x* = 36 chromosomes, comprising of two subgenomes, A and B (Ward 2000; Palomino et al. 2008; Yangquanwei et al. 2013; Kolano et al. 2016; Heitkam et al. 2020). The complexity of the genome due to allotetraploidy and the genetic heterogeneity due to partial outcrossing resulting from the presence of both hermaphroditic and female flowers on the same plant have hampered molecular analysis of quinoa (Maughan et al. 2004; Christensen et al. 2007). Our collaborative group (Yasui et al. 2016) and three subsequent independent groups (Jarvis et al. 2017; Zou et al. 2017; Bodrug-Schepers et al. 2021) have reported the draft genome sequences of quinoa. We generated approximately 140 genotyped quinoa inbred lines suitable for molecular analysis and demonstrated genotype–phenotype relationships among the inbred lines for salt tolerance and key growth traits (Mizuno et al. 2020). We also showed that the quinoa inbred lines can be divided into three genetic subpopulations: the northern highland, southern highland, and lowland (Mizuno et al. 2020). Recently, high-quality chromosome-level genome assemblies have been reported by our collaborating groups for the northern and southern highland lines, J075 and J100, respectively (Kobayashi et al. 2024), and by another independent group for the lowland line QQ74 (Rey et al. 2023). Furthermore, we have developed a technique to analyze the function of endogenous genes in quinoa using virus-induced gene silencing (VIGS) and virus-mediated overexpression (VOX), thereby advancing the field of functional genomics analysis (Ogata et al. 2021).

In the present study, we focused on young quinoa seedlings lacking epidermal bladder cells and examined the physiological responses of all three genotypic lines such as lowland, northern highland, and southern highland lines to high salinity (600 mM NaCl). To further explore the mechanism of salt tolerance in quinoa, we also performed transcriptome analysis, genomic analysis using 18 resequencing data, and VIGS analysis of transporter genes in quinoa plants. We show that CqHKT1 and CqSOS1 are involved in the genotype-dependent Na^+^ exclusion in quinoa, providing insight into salt tolerance mechanisms in quinoa.

## Results

### Quinoa model line Kd thrives in 600 mM NaCl and accumulates Na^+^ in the aerial parts

First, we investigated which growth stage of quinoa should be used for the salt stress test. Epidermal bladder cells on the surface of aerial parts of quinoa are one of the most striking morphological features of quinoa, and their role in salt stress tolerance has been extensively studied and debated (Adolf et al. 2013; Kiani-Pouya et al. 2017; Böhm et al. 2018; Kiani-Pouya et al. 2019; Imamura et al. 2020; Bazihizina et al. 2022; Moog et al. 2023). Recent studies support the notion that epidermal bladder cells are not significantly involved in the salt tolerance of quinoa (Moog et al. 2022). However, given the various possibilities, and with an emphasis on understanding the mechanisms of salt tolerance that are not affected by epidermal bladder cells, this study focused on the mechanisms of salt tolerance in quinoa plants at the young seedling stage (Fig. 1A), which lacks epidermal bladder cells. In the quinoa plants examined in this study, no epidermal bladder cells were present on the cotyledons (Fig. 1B), but epidermal bladder cells form on the first true leaves and all subsequent leaves that emerge (Fig. 1C).

**Figure 1.**
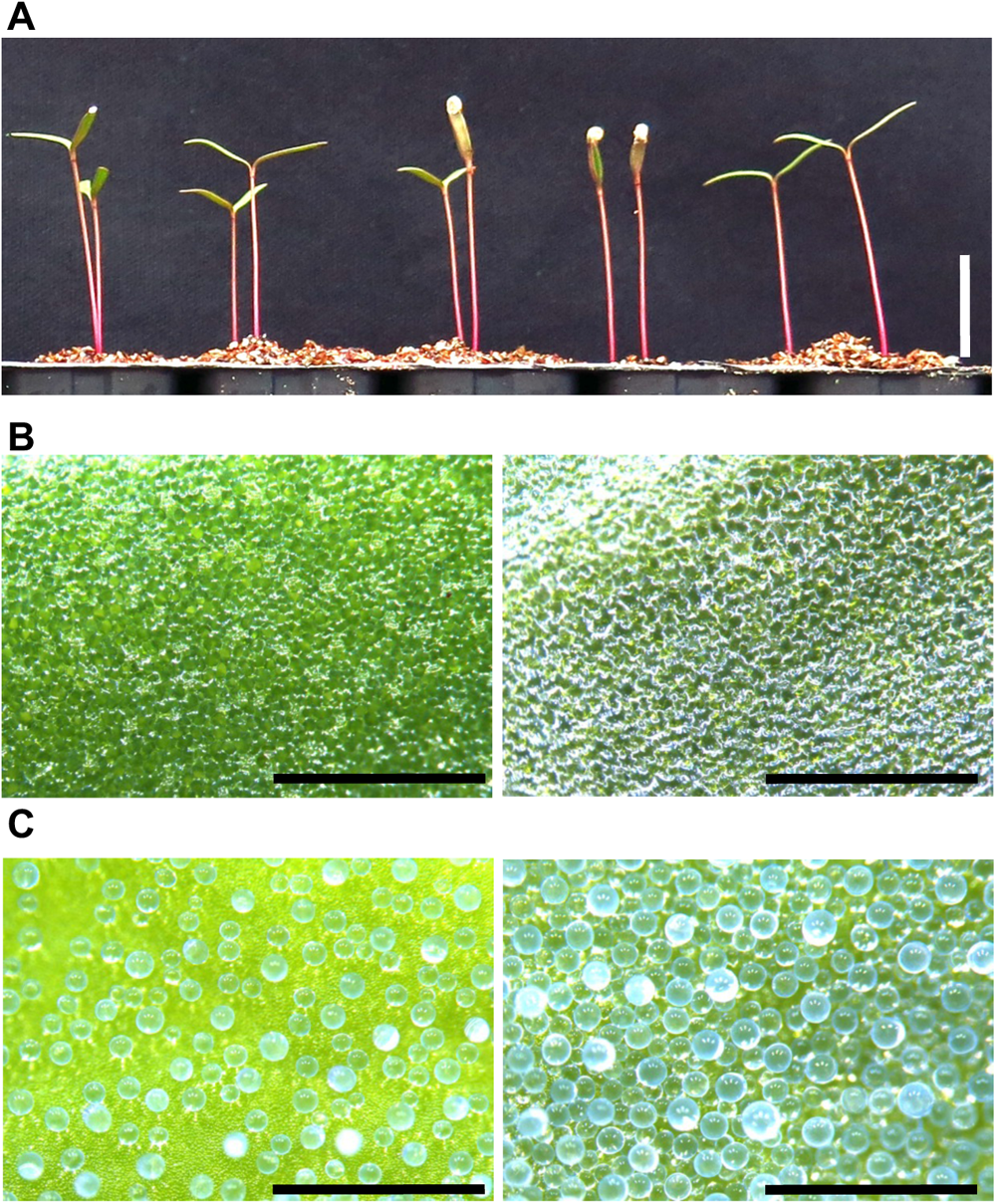
The cotyledons of young quinoa seedlings lack epidermal bladder cells. (A) Young seedlings of the Kd line, a representative model line of quinoa, were grown on vermiculite with RO water for 10 days. Bar, 1 cm. Epidermal surface of cotyledon (B) and developing true leaf (C) were observed by a stereomicroscope. Left and right panels indicate adaxial and abaxial surfaces, respectively. Bars, 1 mm.

Another important consideration is the concentration of NaCl used in salt stress tests. Our previous salt stress tests on quinoa germination have yielded the following results (Mizuno et al. 2020). In the test treated with 300 mM NaCl for 4 days, there was no difference in the results among the lowland, northern highland, and southern highland quinoa lines, but in the test treated with 600 mM NaCl (equivalent to seawater) for 14 days, most of the northern highland lines did not germinate, but the lowland and southern highland lines did, indicating that the results of the 600 mM NaCl-treated trials were genotype related (Mizuno et al. 2020). Interestingly, there was no correlation between the results of the 300 and 600 mM NaCl germination tests (*R*^2^ = 0.01) (Mizuno et al. 2020), suggesting that higher levels of salt stress, such as 600 mM NaCl, are more likely to produce genotype-related responses than are low or moderate levels of salt stress. We therefore used 600 mM NaCl as the salt stress concentration in this study.

We determined the dry weight and Na^+^ and K^+^ accumulations of various tissues of quinoa seedlings up to 4 days after treatment with or without 600 mM NaCl. In this analysis, we used Kd, a sequenced model quinoa line that exhibits uniform growth among individuals and is suitable for molecular biology experiments (Yasui et al. 2016; Mizuno et al. 2020; Ogata et al. 2021) (Fig. 2). Notably, salt treatment did not appear to inhibit growth at most time points up to after 96 h of the salt treatment (Fig. 2, A to C). For all plants examined, the dry weight of the cotyledons increased in a time-dependent manner, whereas the dry weight of the hypocotyls and roots remained mostly unchanged (Fig. 2, A to C). In the salt-treated plants, the cotyledons and hypocotyls showed a time-dependent increase in Na^+^ accumulation, whereas there was no increase in Na^+^ accumulation in control plants (Fig. 2, D and E). In the salt-treated plants, the Na^+^ accumulation in the roots plateaued after 24 h of salt treatment, all but the earliest time points was higher than in the control plants (Fig. 2F). No difference in K^+^ accumulation was observed between the cotyledons of the salt-treated and control plants (Fig. 2G); however, the K^+^ accumulation was lower in the roots of the salt-treated plants than in those of the control plants after 2 h of salt treatment (Fig. 2I). By contrast, K^+^ accumulation in the hypocotyls increased in salt-treated plants but decreased in control plants up to after 6 h of salt treatment (Fig. 2H). After from 12 to 48 h of the salt treatment, there was no difference in K^+^ accumulation between the salt-treated and control plants, but after this time point, K^+^ increased in the control seedlings and decreased in the salt-treated plants (Fig. 2H). The hypocotyls of salt-treated plants may have had a transient increase in potassium levels due to the supply of high-salinity water, while the hypocotyls of untreated control plants may have had a transient decrease in potassium levels due to the supply of water. Thus, the growth of the model quinoa line Kd was not affected by treatment with 600 mM NaCl, but high salinity resulted in the accumulation of Na^+^, mainly in the aerial parts of the plants.

**Figure 2.**
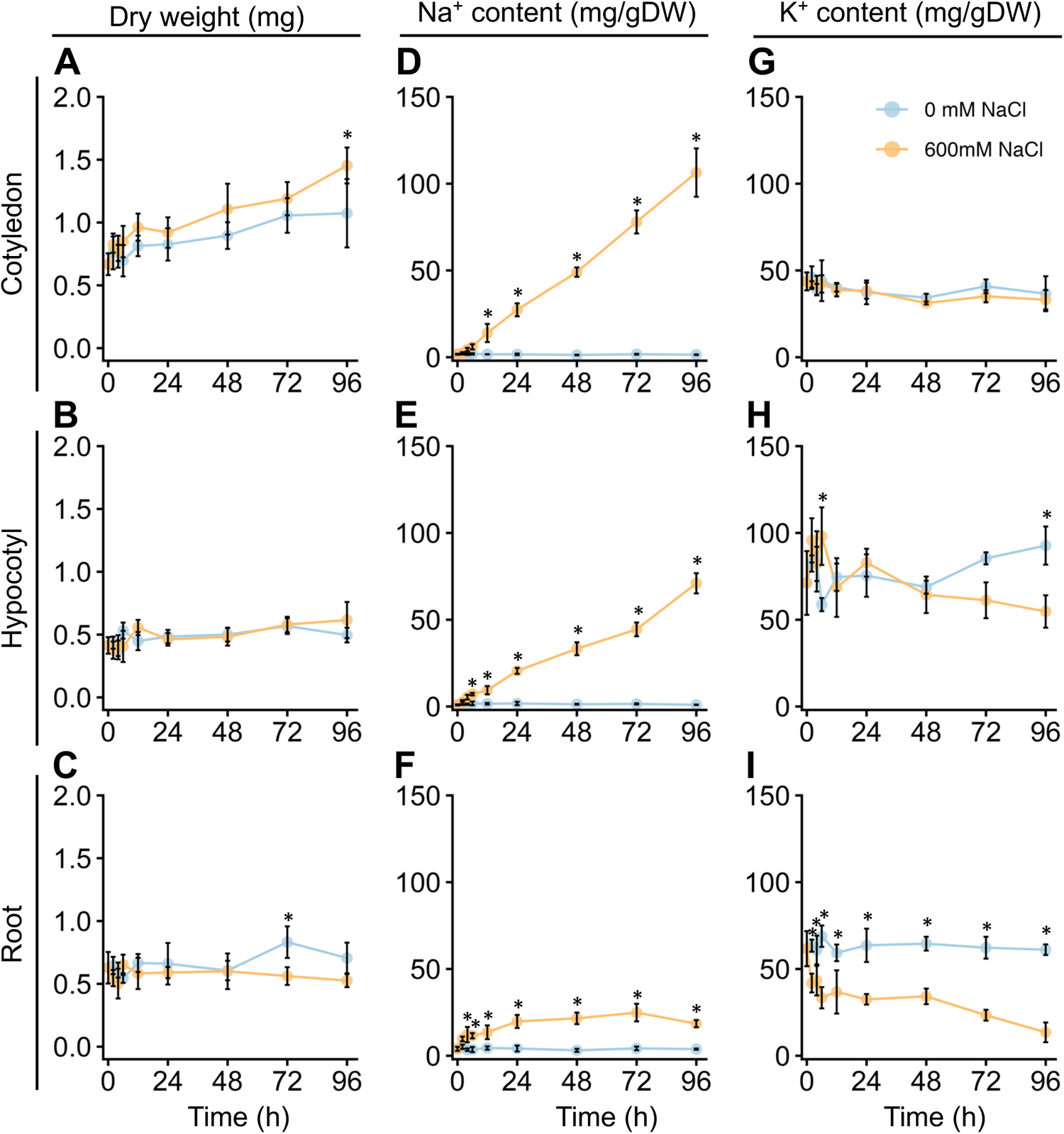
Growth and Na^+^ and K^+^ contents of young quinoa seedlings in response to high salt stress over time. Ten-day-old quinoa seedlings grown on a vermiculite culture system were treated with 0 mM or 600 mM NaCl for 96 h. Cotyledon (top), hypocotyl (middle), and root (bottom) tissues were collected at 0, 2, 4, 6, 12, 24, 48, 72, and 96 h after the salt or control treatment. Dry weight (A, B, C) and Na^+^ (D, E, F) and K^+^ (G, H, I) contents were measured in each tissue. Error bars indicate SD (*n* = 5). \**P* < 0.05, one-way analysis of variance with Tukey’s HSD test was used to evaluate differences between seedlings subjected to the 600 mM NaCl treatment versus the 0 mM NaCl treatment at each time point.

### High salinity does not affect the early seedling growth of all three quinoa genotypes

Previously, we showed that the population of quinoa inbred lines can be genetically divided into three subpopulations: northern highland, southern highland, and lowland subpopulations (Mizuno et al. 2020). Next, we investigated the physiological responses of all three subpopulations, as well as the quinoa model line Kd, which belongs to the lowland subpopulation, to high seawater salinity. Based on the results of our previous genetic structure analysis of quinoa inbred lines (Mizuno et al. 2020), we selected six inbred lines containing 100% of the genetic background of each subpopulation from each subpopulation for this experiment: in addition to Kd, we selected J028, J045, J079, J082, and J122 as representative lines of the lowland subpopulation; J064, J071, J072, J073, J074, and J075 as representative lines of the northern highland subpopulation; J054, J094, J096, J099, J100, and J128 as representative lines of the southern highland subpopulation (Supplementary Table S1). For all three genotypes examined, there was no significant difference in the dry weights of cotyledons, hypocotyls, and roots between seedlings treated or not with 600 mM NaCl at either 24 or 48 h after the salt treatment (Figs. 3 and 4; Supplementary Fig. S1), indicating that this treatement did not affect the initial growth of quinoa seedlings of any of the three genotypes. Interestingly, although the 600 mM NaCl treatment did not affect the growth of young Kd seedlings, it caused a rapid upward (hyponastic) leaf movement (Fig. 3), suggesting that these plants have a physiological response to salt stress.

**Figure 3.**
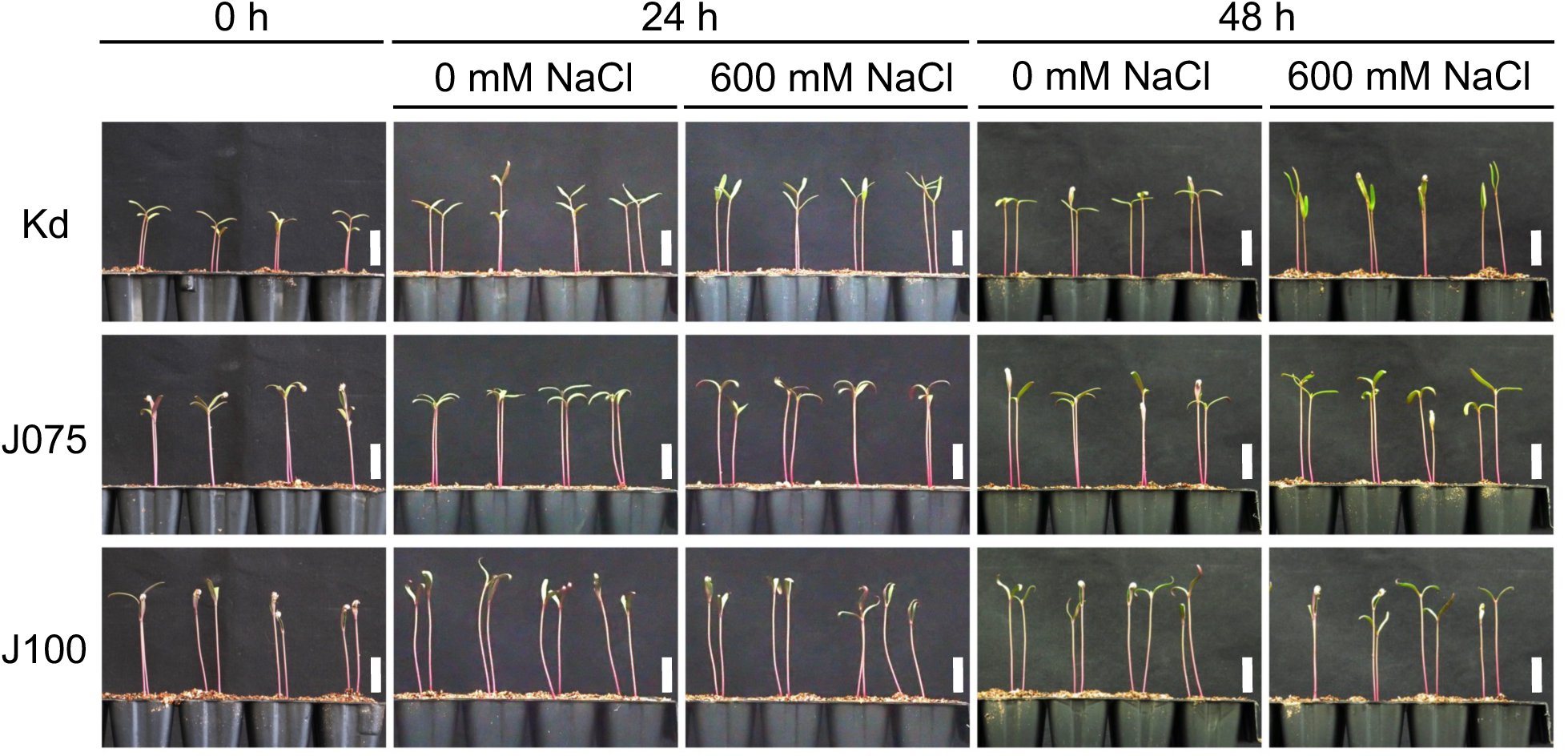
Growth of representative quinoa inbred lines of each genotype during salt stress test. Ten-day-old seedlings of quinoa inbred lines were treated with 0 mM or 600 mM NaCl for 0, 24 and 48 h. Photographs showing representative growth of Kd, a representative inbred line of lowland quinoa, J075, a representative inbred line of northern highland quinoa, and J100, a representative inbred line of southern highland quinoa. Lowland quinoa inbred line Kd, treated with 600 mM NaCl for 24 and 48 h, exhibited rapid upward (hyponastic) leaf movement not seen in the other lines. In some seedlings, especially in the southern highland line J100, the cotyledons are not fully open because the seed coat is still present. Bars, 1 cm.

### Quinoa genotype determines aboveground uptake of Na^+^

We then examined tissue-specific Na^+^ accumulation in young quinoa seedlings treated or not with 600 mM NaCl for 24 and 48 h using six representative inbred lines of each of the three genotypes (Fig. 5A; Supplementary Fig. S2; Supplementary Table S1). In roots, salt-treated plants accumulated 1.4-to 6.2-fold more Na^+^ than controls for all genotypes, while in cotyledons and hypocotyls, salt-treated plants accumulated 7.5-to 67.4-fold more Na^+^ than controls (Fig. 5A; Supplementary Fig. S2). In all salt-treated lines, Na^+^ accumulation tended to be highest in the cotyledons, at intermediate levels in the hypocotyls, and lowest in the roots (Fig. 5A), indicating that young quinoa seedlings grown in a high-salinity environment accumulate more Na^+^ in the aerial parts than in the roots (Fig. 5A; Supplementary Fig. S2). Aboveground Na^+^ accumulation, especially in the cotyledons, tended to be higher in the lowland lines and lower in the southern highland lines (Fig. 5A).

**Figure 4.**
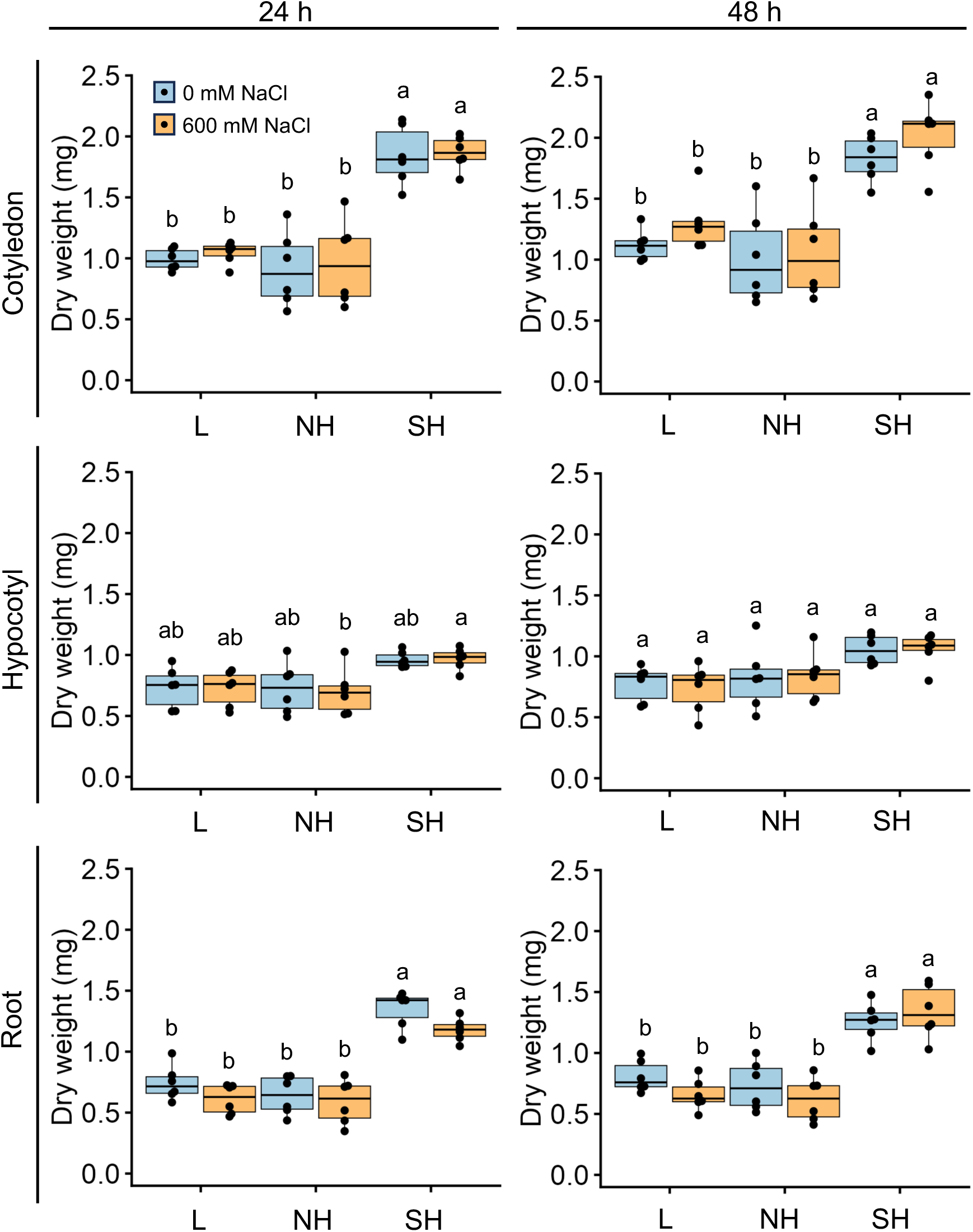
Genotype-specific growth in response to high salt stress over time. Ten-day-old seedlings of quinoa inbred lines were treated with 0 or 600 mM NaCl for 24 h (left) and 48 hours (right). Data for lowland (L) lines include Kd, J028, J045, J079, J082, and J122; data for northern highland (NH) lines include J064, J071, J072, J073, J074, and J075; data for southern highland (SH) lines include J054, J094, J096, J099, J100, and J128 (Supplementary Table S1). Average dry weight of cotyledons, hypocotyls, and roots of each line is indicated by dots in the box plot. Different letters indicate significant differences among subpopulations by Tukey’s HSD test (*P* < 0.05). Dry weight data for individual lines are shown in Supplementary Fig. S1.

**Figure 5.**
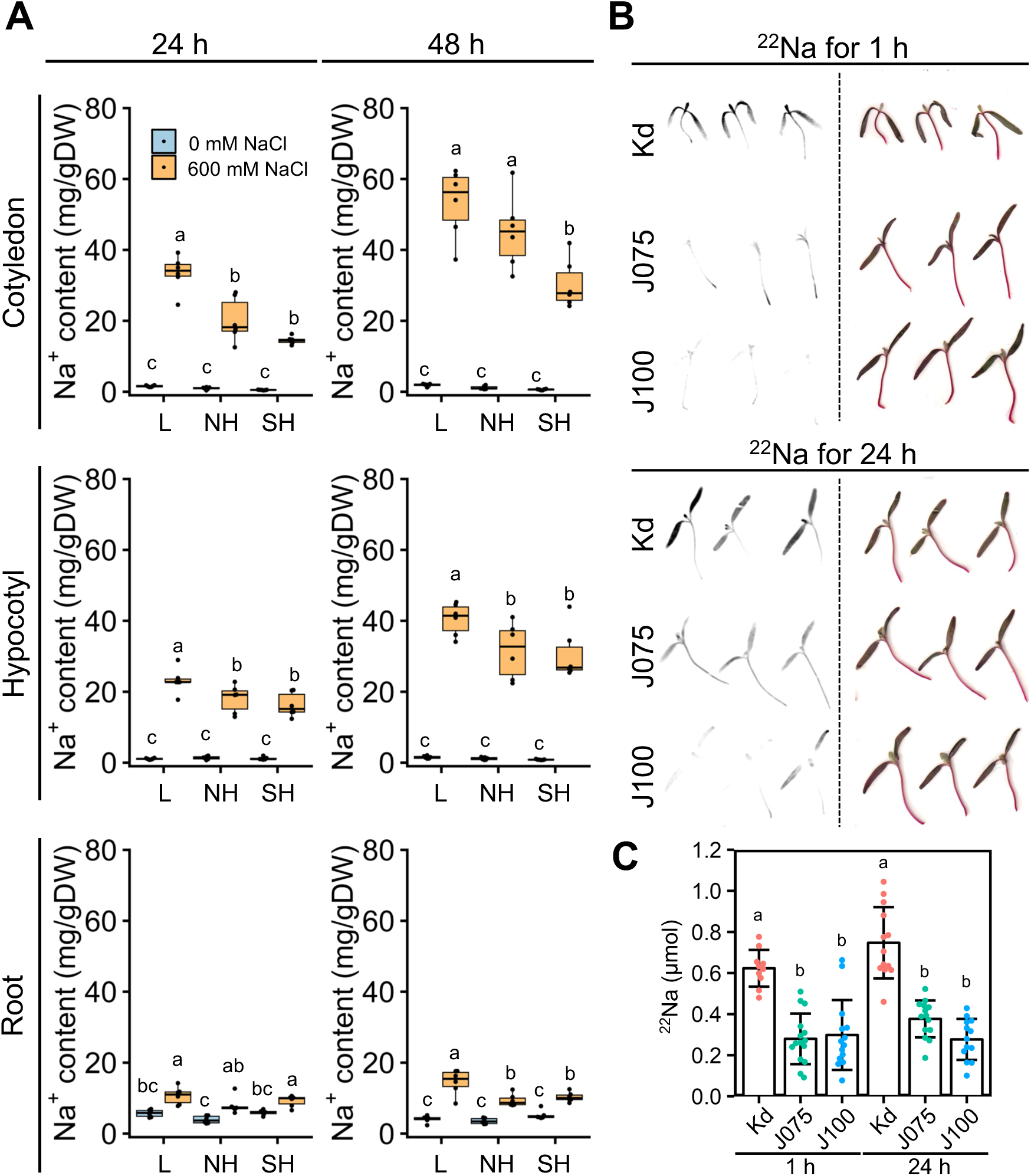
Genotype-specific Na^+^ accumulation and uptake in response to high salt stress over time. (A) Ten-day-old seedlings of quinoa inbred lines were treated with 0 or 600 mM NaCl for 24 h (left) and 48 h (right). Data for lowland (L) lines include Kd, J028, J045, J079, J082, and J122; data for northern highland (NH) lines include J064, J071, J072, J073, J074, and J075; data for southern highland (SH) lines include J054, J094, J096, J099, J100, and J128 (Supplementary Table S1). Average Na^+^ content in cotyledons, hypocotyls, and roots of each line is indicated by dots in the box plot. Different letters indicate significant differences among subpopulations by Tukey’s HSD test (*P* < 0.05). Na^+^ accumulation data for individual lines are shown in Supplementary Fig. S2. (B) Autoradiography (left) and actual photographs (right) of above-ground parts of quinoa Kd, J075, and J100 seedlings radiolabeled with ^22^Na. The seedlings were soaked in water containing ^22^Na for 1 and 24 hours. (C) ^22^Na content of the aerial part of each quinoa line, calculated from a photostimulated luminescence value and a calibration curve based on the spots (*n* = 10–15 seedlings). Different letters indicate significant differences among inbred lines by Tukey’s HSD test (*P* < 0.05).

To determine whether this difference in aboveground Na^+^ accumulation by genotype was due to differences in growth or Na^+^ uptake by genotype, we performed uptake experiments with radiolabeled ^22^Na using Kd as a representative lowland line, J075 as a representative northern highland line, and J100 as a representative southern highland line (Fig. 5B). At 1 and 24 h after transferring the young seedlings to the ^22^Na-containing medium, we observed significantly stronger ^22^Na signals in the aboveground parts of the lowland Kd seedlings than in those of J075 and J100 (Figs 5, B and C). Thus, Na^+^ uptake in the aerial parts of quinoa is higher in the lowland line and lower in the northern highland and southern highland lines, demonstrating that Na^+^ uptake in the aerial parts of quinoa is defined by the genotype.

### High salinity reduces root K^+^ accumulation in all three genotypes

We further assessed tissue-specific K^+^ accumulation in young quinoa seedlings treated or not with 600 mM NaCl for 24 and 48 h using six representative inbred lines of each of the three genotypes (Fig. 6; Supplementary Fig. S3; Supplementary Table S1). In cotyledons and hypocotyls, salt treatment tended to slightly reduce K^+^ accumulation in many of the lines (Fig. 6). Regardless of the salinity conditions, K^+^ accumulation tended to be slightly lower in the aerial parts of southern highland lines than in those of the lowland and northern highland lines (Fig. 6). By contrast, in roots, salt treatment significantly reduced K^+^ accumulation in all lines examined, and K^+^ accumulation differed little among genotypes (Fig. 6). The results obtained so far showed that Na^+^ accumulation in the cotyledons of the salt-treated young seedlings is different among the genotypes but that there is no clear difference in Na^+^ and K^+^ accumulation in the roots among the genotypes (Figs. 5 and 6). We therefore concluded that salt treatment generally increases Na^+^ accumulation and decreases K^+^ accumulation in all tissues across all genotypes (Figs. 5 and 6).

**Figure 6.**
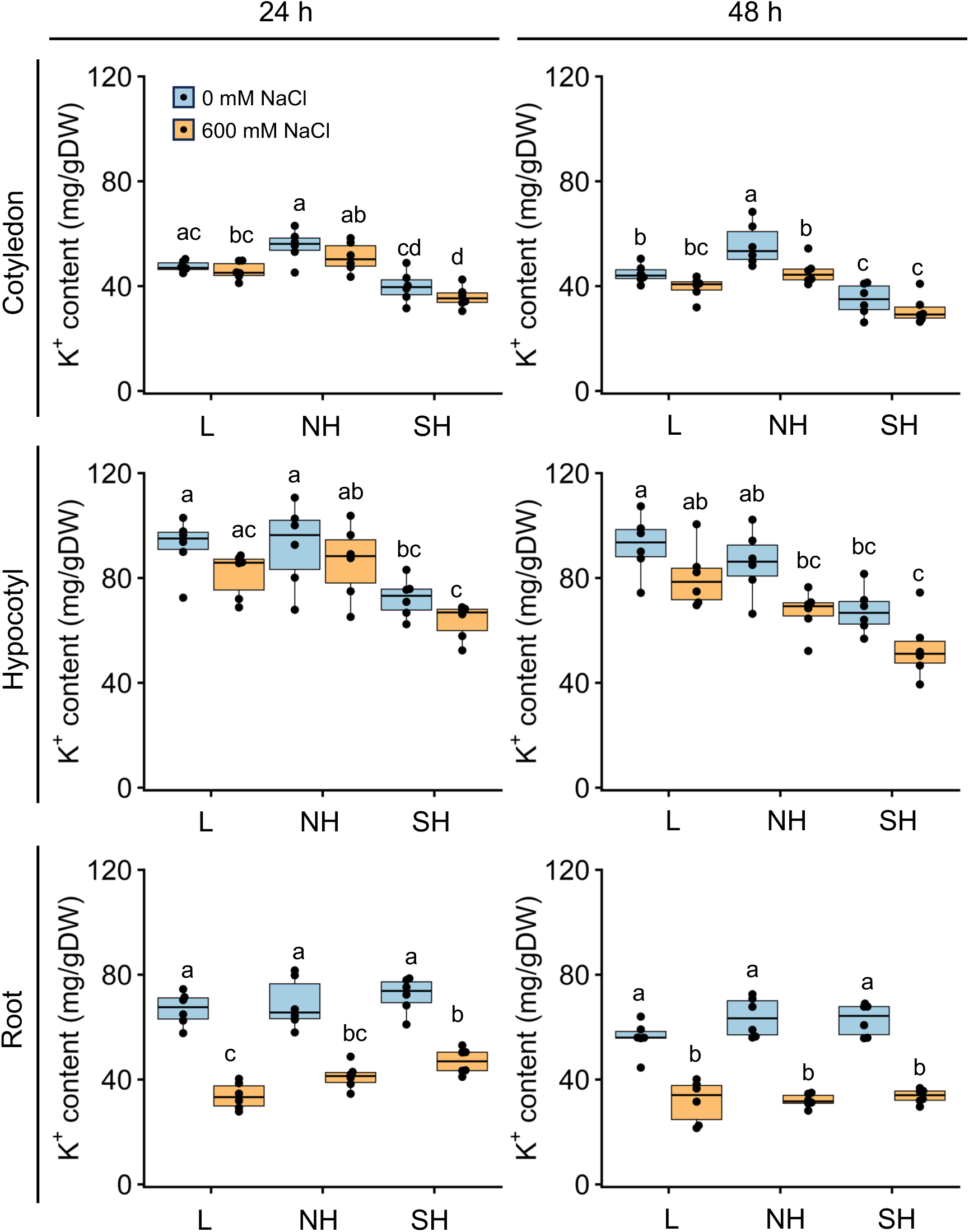
Genotype-specific K^+^ accumulation in response to high salt stress over time. Ten-day-old seedlings of quinoa inbred lines were treated with 0 or 600 mM NaCl for 24 h (left) and 48 h (right). Data for lowland (L) lines include Kd, J028, J045, J079, J082, and J122; data for northern highland (NH) lines include J064, J071, J072, J073, J074, and J075; data for southern highland (SH) lines include J054, J094, J096, J099, J100, and J128 (Supplementary Table S1). Average K^+^ content in cotyledons, hypocotyls, and roots of each line is indicated by dots in the box plot. Different letters indicate significant differences among subpopulations by Tukey’s HSD test (*P* < 0.05). K^+^ accumulation data for individual lines are shown in Supplementary Fig. S3.

### Gene expression profile in response to high salt stress varies widely among genotypes

To investigate the effect of high salt stress on gene expression in young quinoa seedlings, we analyzed the gene expression profiles of cotyledons and roots after 0, 6, and 24 h of treatment with or without 600 mM NaCl (Fig. 7). We used Kd as a representative lowland line, J075 as a representative northern highland line, and J100 as a representative southern highland line (Fig. 7). In the RNA-seq experiment, we identified 8,033 and 4,121 differentially expressed genes (DEGs) (|log_2_(fold change (FC))| ≥ 1, transcripts per million (TPM) value > 0, false discovery rate (FDR) < 0.05) in response to high salt stress in at least one sampling in cotyledons and roots of each line, respectively (Fig. 7; Supplementary Table S2). We then performed hierarchical clustering and gene ontology (GO) enrichment analysis of these genes. In cotyledons, one gene set, designated cluster A, was enriched in genes involved in biological processes related to cell wall organization (Fig. 7; Supplementary Figs. S4; Supplementary Table S3). Two other gene sets, designated clusters B and C, were enriched in genes involved in biological processes related to photosynthesis and cellular response to hypoxia (Fig. 7; Supplementary Figs. S5 and S6; Supplementary Table S3). These results indicate that, in cotyledons of the salt-stressed lowland Kd line, the expression of genes related to cell wall composition is up-regulated, while the expression of genes related to photosynthesis and oxygen deprivation response is down-regulated. In roots, one gene set, designated cluster D, was enriched in genes involved in biological processes related to responses to water deprivation and ABA (Fig. 7; Supplementary Figs. S7; Supplementary Table S4). Two additional gene sets, designated clusters E and F, were enriched in genes involved in biological processes related to cellular response to hypoxia, defense response to bacteria, and hydrogen peroxide catabolic process (Fig. 7; Supplementary Figs. S8 and S9; Supplementary Table S4). These results indicate that all lines tested, regardless of the genotype, showed increased expression of genes involved in responses to water deprivation and ABA in their roots under high salt stress. Interestingly, in the roots of the highland lines J075 and J100, salt stress specifically led to a marked increase in the expression of genes related to the hypoxia response, defense response against bacteria, and production of reactive oxygen species (ROS). These observed differences in gene expression between the roots of the lowland quinoa line Kd, which exhibit striking increases in Na^+^ concentration in the aerial parts following high salt treatment, and southern highland quinoa J100, which exhibit lower increases in Na^+^ concentration in the aerial parts following high salt treatment, are expected to provide clues into the mechanisms underlying the high salt stress response in quinoa.

**Figure 7.**
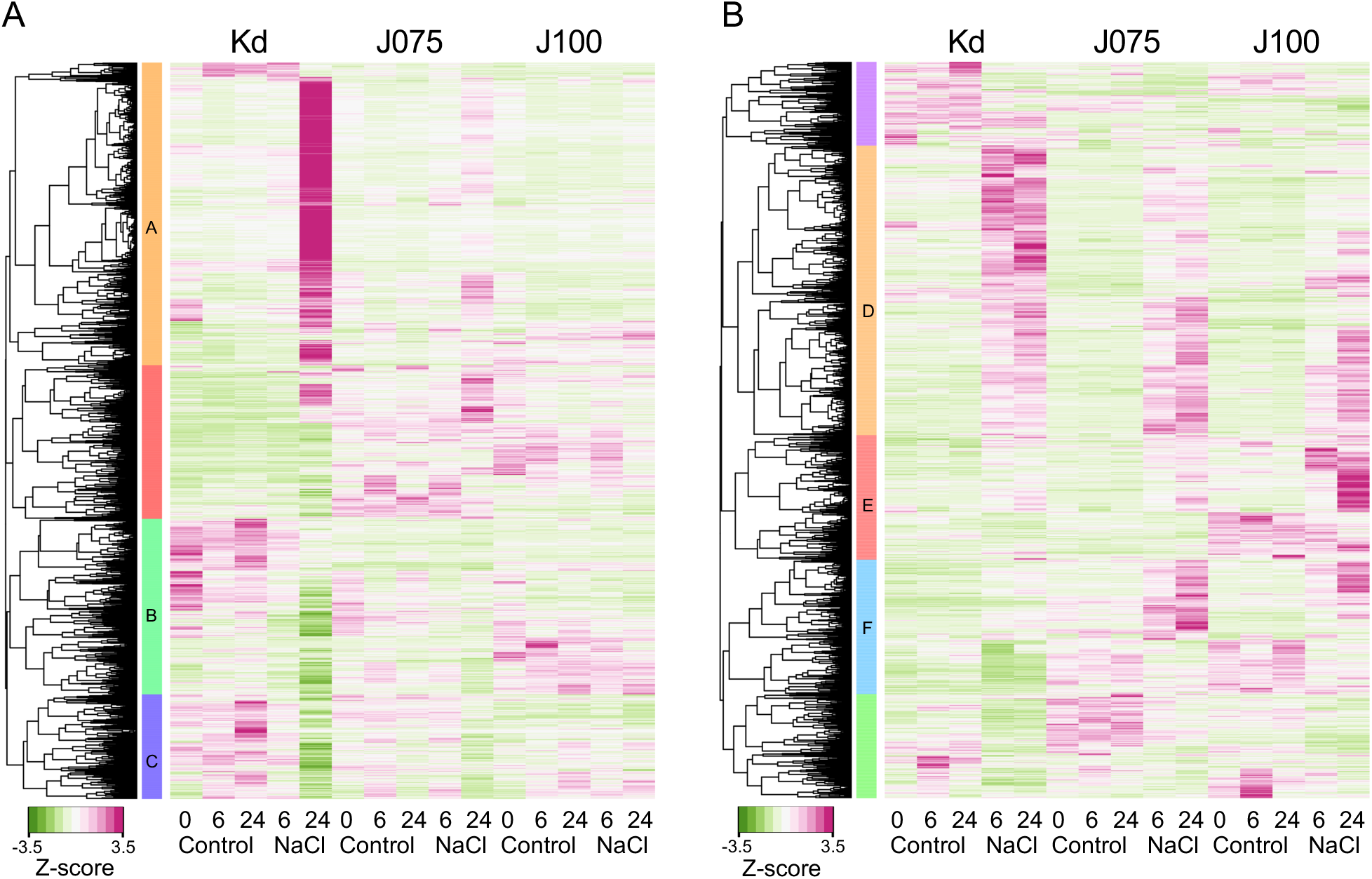
Tissue-specific salt stress–responsive gene expression in genotype representative lines. Hierarchical clustering of 8,303 and 4,121 differentially expressed genes (DEGs) in the RNA-seq experiments (|log2(FC)| ≥ 1, TPM value > 0, FDR < 0.01) that were identified in cotyledons (A) and roots (B) of each genotype representative line subjected to either the 6 or 24 h of NaCl treatment compared to the control (0 mM NaCl treatment) (Supplementary Table S2). Ten young quinoa seedlings were sampled at each time point of each treatment and replicated in triplicate. Six gene sets, designated clusters A through F, of the nine clusters identified in total were enriched in genes involved in biological processes related to characteristic responses (see Supplementary Figs. 4–9 for details). Heat map showing z-scaled TPM of DEGs.

### *CqHKT1* and *CqSOS1* gene expression levels are genotype dependent

So far, we have analyzed gene expression responses to high salt stress, but salt stress–tolerant plants often do not show significant expression of key genes involved in the salt stress response due to constitutive expression of key genes such as *HKT1* and *SOS1* (Taji et al. 2004; Katschnig et al. 2015). HKT1 (Rubio et al. 1995; Horie et al. 2001; Rus et al. 2001) and SOS1 (Shi et al. 2000; Shi et al. 2002) proteins are considered to be among the most important transporters involved in salt stress tolerance in a wide range of plant species (Yang and Guo 2018; Ali et al. 2021). SOS1, a plasma membrane Na^+^/H^+^ antiporter, mediates Na^+^ efflux out of the root and Na^+^ loading into the xylem, while the channel-like protein HKT1 mediates the reverse flux of Na^+^ unloading off the xylem (Yang and Guo 2018; Ali et al. 2021). The quinoa sodium transporter *CqHKT1* gene has been reported to have a root-expressed *CqHKT1;1* and a shoot-expressed *CqHKT1;2* (Böhm et al. 2018). *CqSOS1* is also conserved in the quinoa genome and is mainly expressed in roots (Maughan et al. 2009). The expression of these genes remained unchanged in quinoa subjected to salt stress treatment (Maughan et al. 2009; Böhm et al. 2018). The results of these reports are largely consistent with those of our RNA-seq analysis (Supplementary Fig. S10).

To explore the genetic variation in the quinoa inbred lines examined in this study, we employed resequencing to analyze 18 of the quinoa inbred lines: J028, J045, J079, J082, J122, and Kd as representative lowland lines; J064, J071, J072, J073, J074, and J075 as representative northern highland lines; and J054, J094, J096, J099, J100, and J128 as representative southern highland lines (Supplementary Table S1). Genomic analysis showed that the *CqHKT1;1* gene in lines J072, J073, and J075 had a single nucleotide insertion in the first exon, which shifted the reading frame (Supplementary Fig. S11). Expression of this disrupted gene was completely undetectable (Supplementary Fig. S12). In lines J064 and J071, a 6.9-kb deletion was found in the 5′ flanking region of the *CqSOS1* gene (Supplementary Fig. S13), and almost no gene expression was observed (Supplementary Fig. S12). By contrast, the other lines examined showed full-length gene expression with no mutations in or near the *CqHKT1;1* or *CqSOS1* genes (Supplementary Figs. S14 to S16).

Reverse transcription quantitative PCR (RT-qPCR) analysis revealed that *CqHKT1;2* in cotyledons and *CqHKT1;1* and *CqSOS1* in roots did not show significant changes in gene expression in response to salt stress in many of the lines examined, but there were significant differences in expression levels among genotypes with and without salt stress treatment (Fig. 8; Supplementary Fig. S12). However, the genotype dependence of the expression levels of these genes was higher in *CqHKT1;2* and *CqSOS1* and tended to be lower in *CqHKT1;1* (Fig. 8; Supplementary Fig. S12). In most cases, *CqHKT1;1* and *CqHKT1;2* were more strongly expressed in the lowland lines than in the highland lines, whereas *CqSOS1* was more strongly expressed in the southern and northern highland lines than in the lowland lines (Fig. 8; Supplementary Fig. S12). In addition, many sequence polymorphisms were observed around the promoter regions of the *CqHKT1;1*, *CqHKT1;2*, and *CqSOS1* genes between lowland and highland quinoa lines (Supplementary Figs. S14 to S16), strengthening the hypothesis that the expression levels of these transporter genes are genotype dependent. Given that natural gene expression levels of transporters are associated with Na^+^ accumulation and salt tolerance (Oh et al. 2009; Jha et al. 2010), these results support the notion that the function and activity of the transporters are defined by genotype in quinoa.

**Figure 8.**
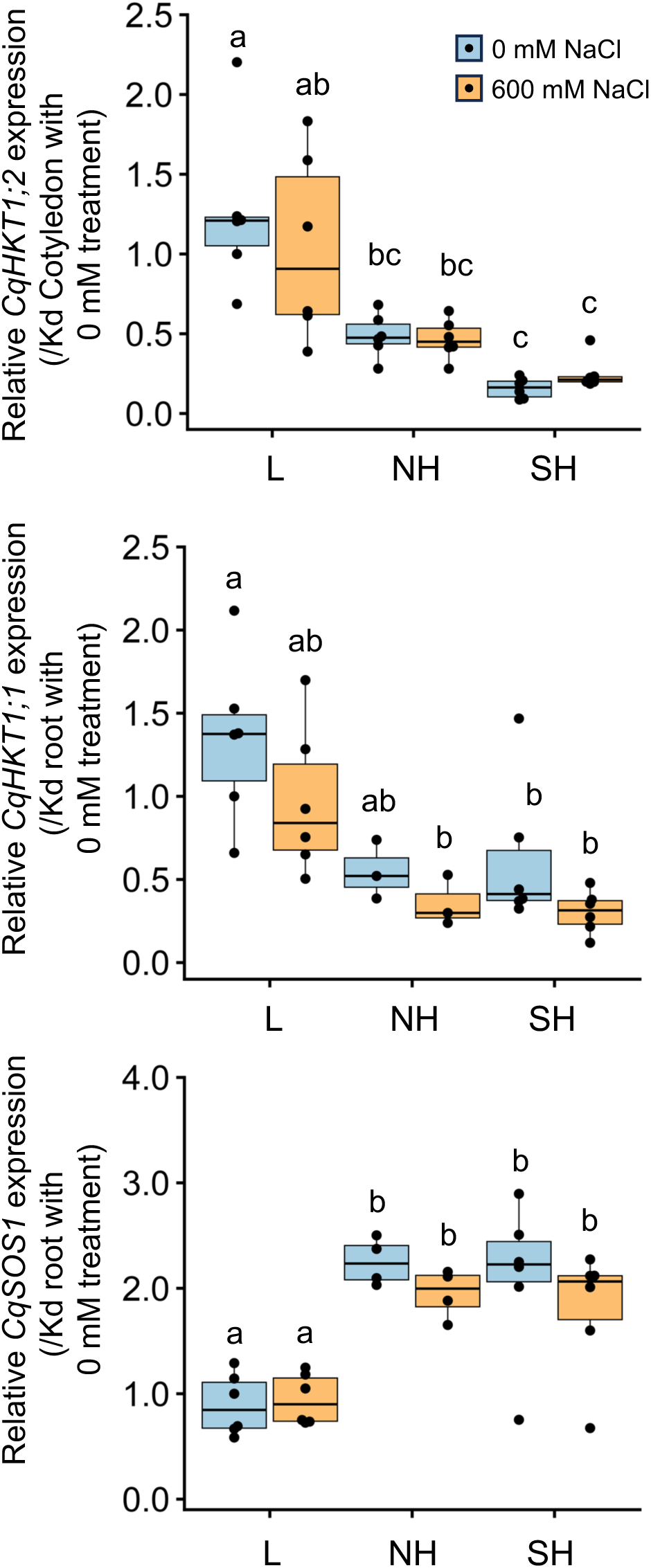
Genotype-specific expression of transporter genes in response to high salt stress. Ten-day-old seedlings of quinoa inbred lines were treated with 0 or 600 mM NaCl for 24 h. Gene expression in cotyledons was examined for *CqHKT1;2*, which is mainly expressed in cotyledons, and gene expression in roots was examined for *CqHKT1;1* and *CqSOS1*, which are mainly expressed in roots. The transcript levels of *CqHKT1;2*, *CqHKT1;1*, and *CqSOS1* genes were normalized to those of *CqUBQ10* as an internal control gene. Average expression levels of CqHKT1;1, CqHKT1;2, and CqSOS1 are shown as dots in the box plots relative to the gene expression levels in each corresponding tissue in the non-salt-treated young Kd seedlings. Data for lowland (L) lines include Kd, J028, J045, J079, J082, and J122; data for northern highland (NH) lines include J064, J071, J072, J073, J074, and J075; and data for southern highland (SH) lines include J054, J094, J096, J099, J100, and J128 (Supplementary Table S1). The *CqHKT1;1* genes in J072, J073, and J075 and the *CqSOS1* genes in J064 and J071 had an adenine insertion in the first exon and a 6.9-kb deletion in the 5′ region of the gene’s coding region and the corresponding transcripts were not full-length or functional; gene expression data for these genes are not included. Different letters indicate significant differences among subpopulations based on a Tukey’s HSD test (*P* < 0.05). Relative expression levels for individual lines are shown in Supplementary Fig. S12.

### CqHKT1 and CqSOS1 mediate Na^+^ exclusion in quinoa

To determine whether the CqHKT1 and CqSOS1 transporters are involved in Na^+^ transport in quinoa, we suppressed the gene expression of these transporters using the VIGS method with an ALSV vector, the only method currently available to analyze endogenous gene function in quinoa (Ogata et al. 2021) (Fig. 9). Because this ALSV-VIGS experimental system requires virus inoculation of leaves with epidermal bladder cells and sampling of uninoculated upper leaves with epidermal bladder cells, the transport ability of the transporters was analyzed in older quinoa plants, which have true leaves with epidermal bladder cells, rather than in seedlings, which lack epidermal bladder cells. In addition, a model quinoa line, the lowland line Kd, was used in this experiment. This is because lowland lines tend to accumulate more Na^+^ in the aerial parts under high salinity conditions, unlike southern highland lines, and are therefore more suitable for analyzing the function of these transporters. RT-qPCR analysis revealed that *CqHKT1;1*, *CqHKT1;2*, and *CqSOS1* expression was significantly down-regulated in the uninoculated upper leaves of the plants inoculated with ALSV-CqHKT1;1, ALSV-CqHKT1;2, and ALSV-CqSOS1, respectively, compared with those inoculated with ALSV-WT in each experiment (Fig. 9B). Knockdown of *CqHKT1;1*, *CqHKT1;2*, and *CqSOS1* had no significant effect on plant height (Fig. 9, B and C) but increased the Na^+^ content in the uninoculated upper leaves of salt-treated plants by 56%, 47%, and 55%, respectively, indicating that reduced expression of these transporter genes results in increased salt accumulation in plants (Fig. 9D). Considering that these transporters generally function to suppress Na^+^ transport to leaves in a variety of plants (Apse and Blumwald 2007; Plett and Møller 2010), these results demonstrate that *CqHKT1;1*, *CqHKT1;2*, and *CqSOS1* function in Na^+^ exclusion under high-salinity conditions in quinoa.

**Figure 9.**
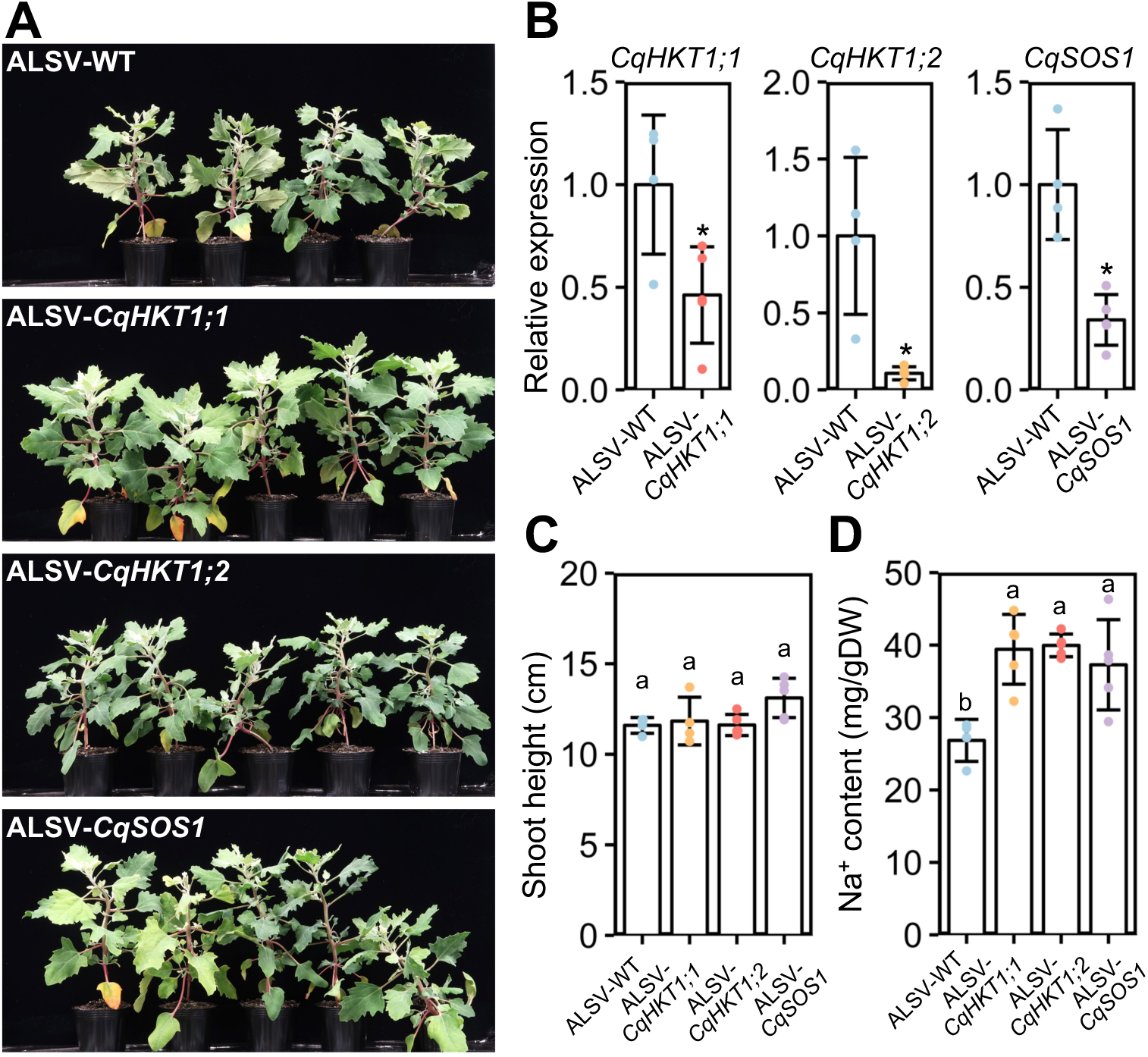
Knockdown of *CqHKT1;1*, *CqHKT1;2* or *CqSOS1* increased Na^+^ content in the uninoculated upper leaves of salt-treated plants. (A) Representative images of plants inoculated with ALSV-WT, ALSV-CqHKT1;1, ALSV-CqHKT1;2, or ALSV-CqSOS1 for 10 days and then treated with 300 mM NaCl for 21 days. (B) RT-qPCR quantification of *CqHKT1;1, CqHKT1;2* and *CqSOS1* transcripts in the uninoculated upper leaves of plants inoculated with the indicated inocula. Data were normalized to the *CqUBQ10* levels and are presented as means ± SD (*n* = 4 for ALSV-WT; *n* = 5 for ALSV-each target gene). (C) Plants were treated with 300 mM NaCl 10 days after inoculation, and shoot height was measured 21 days later. (D) Na^+^ content of the lowest uninoculated upper leaves of virus vector inoculated plants on day 21 of NaCl treatment. Data show mean ± SD (*n* = 4 in ALSV-WT and *n* = 5 in ALSV-each target gene). Asterisks and different lowercase letters indicate significant difference at *P* < 0.05, as determined using Student’s *t*-test and Tukey’s HSD test.

## Discussion

Using our previously reported VIGS application with the ALSV vector (Ogata et al., 2021), we show for the first time that the quinoa transporters CqHKT1;1, CqHKT1;2, and CqSOS1 contribute to Na^+^ exclusion in quinoa plants (Fig. 9). Considering that previous findings of transporters involved in salt transport in quinoa were based solely on genomic and gene expression analysis (Zou et al. 2017; Shi and Gu 2020), the demonstration of transporter function in quinoa in this study is a major step towards understanding the salt tolerance mechanism in quinoa. However, much remains to be resolved. There are no significant differences in Na^+^ or K^+^ accumulation or growth between the roots of northern highland lines that harbor or lack functional *CqHKT1;1* or *CqSOS1* genes (Figs. 3 to 6; Supplementary Figs. S1 to S3). By contrast, no northern highland lines lack functional versions of both *CqHKT1;1* and *CqSOS1* functioning in the roots (Supplementary Figs. S11 to S13). These observations suggest that, in northern highland quinoa lines, it may be sufficient to have either *CqHKT1;1* or *CqSOS1*, or that other genes may be involved in Na^+^ exclusion in leaves. In leaves, the inward rectifier CqHKT1;2 has been reported to play a key role in Na^+^ loading of bladder cells under salt stress (Böhm et al. 2018), but its primary role in quinoa needs to be clarified in the context of findings from homologous genes in other plants. In this study, we show that both CqHKT1;2, which functions mainly in leaves, and CqHKT1;1 and CqSOS1, which function mainly in roots, are involved in Na^+^ exclusion through gene repression (Supplementary Fig. S10). The roles of the transporters involved in salt tolerance in quinoa and the properties that make these transporters superior to those of other plants that are less salt tolerant than quinoa remain to be elucidated. Thus, the present study provides a basis for solving these important mysteries of the salt tolerance mechanism.

We found that young quinoa seedlings at a growth stage lacking epidermal bladder cells grow as well as non-salt-treated control plants under 600 mM NaCl (Figs. 2 to 4; Supplementary Fig. S1), indicating that the salt tolerance of young quinoa seedlings is far greater than that of model and crop plants (Nanjo et al. 1999; Zapata et al. 2017). Given that the Na^+^ influx into plant cells cause K^+^ efflux, a phenomenon involved in many physiological processes, disrupting K^+^ homeostasis (Wang et al. 2009; Demidchik 2014; Pottosin and Shabala 2014; Assaha et al. 2017), the ability of quinoa to maintain high K^+^ levels even under high-salinity conditions may contribute to its high salt tolerance (Fig. 6). In fact, aboveground K^+^ contents in quinoa seedlings do not change significantly under high-salt conditions (Fig. 6). More interestingly, the differences among quinoa genotypes in K^+^ accumulation under high-salt conditions are smaller than those in Na^+^ accumulation (Figs. 5 and 6). In terms of gene expression, it is striking that cell wall–related genes were enriched in Na^+^-accumulating leaves (Fig. 7; Supplementary Figs. S4 to S6; Supplementary Table S3). These findings support previous reports that cell wall remodeling occurs under salt stress in the other plants (Shen et al. 2014; Wang et al. 2019), but how this relates to quinoa’s unique salt tolerance mechanism remains to be elucidated. In roots, the expression of genes associated with the ABA pathway and water deprivation response was up-regulated regardless of the genotype, while the expression of genes associated with the hypoxia response, defense response to biotic stress, and the ROS pathway was up-regulated in the highland lines, especially in the southern highland lines (Fig. 7; Supplementary Figs. S7 to S9; Supplementary Table S4). The results of these GO analyses suggest that both hypoxia and salt stress induce common defense responses, such as Ca^2+^ signaling, membrane polarization, and ROS production (Wang et al. 2017; Cackett et al. 2022), and that these responses may differ between highland and lowland lines. Recent physiological and biochemical findings from five highland quinoa lines suggest that leaf osmotic regulation, K^+^ retention, Na^+^ exclusion, and ion homeostasis are the main physiological mechanisms conferring salt stress tolerance to highland quinoa lines (Cai and Gao 2020); therefore, to elucidate the mechanisms of salt tolerance in quinoa, our focus should turn to ion transport.

We further demonstrated that quinoa genotypes differ significantly in their tendency to accumulate Na^+^ in the aerial parts of the plant under high-salinity stress (Figs 2 and 5; Supplementary Fig. S2). This is the first report on the relationship between genotype and salt accumulation in quinoa using inbred lines based on genomic analysis, and is consistent with a previous observation that, among the lines examined, the genotypes originated from the Bolivian Altiplano had the lowest levels of sodium in leaves under saline conditions (Shabala et al. 2013). Under high-salinity conditions, the lowland quinoa lines accumulated more Na^+^ in the aerial parts, whereas the southern highland lines did not accumulate much Na^+^ in the aerial parts (Figs. 2 and 5; Supplementary Fig. S2). We also showed that the differences in Na^+^ accumulation among these genotypes under high-salinity conditions is due to the differences in the aboveground uptake of Na^+^ between highland and lowland lines under high-salinity conditions (Fig. 5B). In addition, many sequence polymorphisms around the promoter regions of *CqHKT1;1*, *CqHKT1;2*, and *CqSOS1* were observed between lowland and highland quinoa lines (Supplementary Figs. S14 to S16). The lowland and southern highland lines have both root transporters, CqHKT1;1 and CqSOS1, whereas five of the six northern highland lines have only one of the two root transporters (Supplementary Fig. S12), supporting the notion that only functional transporters were transmitted from the northern highlands to the southern highlands and lowlands. These results therefore suggest that the Na^+^ transporter–mediated phenotype involved in quinoa Na^+^ exclusion is closely related to quinoa genotype, consistent with a previous genome-based hypothesis that domestication first occurred in the northern highlands of the Altiplano (Mizuno et al. 2020).

## Materials and Methods

### Plant materials, growth conditions, and salt treatments

The quinoa inbred lines (Yasui et al. 2016; Mizuno et al. 2020; Ogata et al. 2021) used in this study are listed in Supplementary Table S1. For salt stress tests, quinoa seeds were sown and grown in vermiculite-filled cell trays in 8.2-L trays containing reverse osmosis (RO) water for 10 days in a growth chamber set at 22 ± 2°C under a 12-h-light/12-h-dark photoperiod, with light supplied at a 75 ± 25 µmol photons/m^2^/s. Ten days after seeding, these cell trays were transferred to trays containing 0 or 600 mM NaCl RO water. After the salt or control treatment, individual plants were pulled from their cells at the designated time and cut into roots, hypocotyls, and cotyledons for sampling. For VIGS analysis, quinoa Kd seeds were sown in a peat moss mixture (Jiffy Mix, Sakata Seeds, Yokohama, Japan) in a cell tray. After 7 days, the seedlings were transferred to a standard potting mix (Tsuchitaro, Sumitomo Forestry, Tokyo, Japan) in 0.11-L pots and grown in a temperature-controlled phytotron set at 22 ± 2°C under a 12-h-light/12-h-dark photoperiod for 7 days.

### Determination of Na^+^ and K^+^ ions in quinoa seedling tissues

The Na^+^ and K^+^ measurements were performed as previously described (Cataldi et al. 2003), with slight modifications. Cotyledon, hypocotyl, and root tissues of quinoa seedlings were collected after salt stress treatment and oven-dried at 60°C for 2 days. Tissue extracts were prepared by adding 1 mL ultrapure water to ground tissue using a ShakeMaster (BMS, Japan) and 2-mm zirconia beads (1,100 rpm, 2min). The suspensions were heated at 90°C for 20 min and centrifuged at 21,500 *g* for 15 min at 4°C. The ionic contents were analyzed using a high-performance liquid chromatography (HPLC) system (Jasco, Japan, PU-4185 pump, CO-4060 column oven, AS-4150 autosampler) with a CD-200 conductivity detector (Shodex, Japan). Na^+^ and K^+^ were separated using a Shim-pack IC-C4 with Shim-pack IC-C4 guard column (Shimadzu, Japan) and eluted with 2.5 mM oxalic acid. The peak area and concentration of the ions were calculated using ChromNAV Ver.2 software (Jasco, Japan).

### ^22^Na experiment for sodium uptake assay

Quinoa Kd, J100, and J075 seeds were sown on mesh floating in a modified MGRL solution (Kobayashi et al. 2013) containing 0.175 mM sodium phosphate buffer, 0.4 mM NaNO_3_, 0.3 mM KNO_3_, 0.2 mM CaCl_2_, 0.15 mM MgSO_4_, 3 μM H_3_BO_3_, 0.86 μM Fe(III)-EDTA, 1.03 μM MnSO_4_, 0.1 μM CuSO_4_, 0.1 μM ZnSO_4_, 0.24 nM (NH_4_)_6_Mo_7_O_24_ 4H_2_O, 1.3 nM CoCl_2_, and 2 mM MES (pH 5.6). After 6 days, the seedlings were transferred to hydroponic solution containing ^22^Na (1 kBq/mL, Chiyoda Technol Co. Japan). The seedlings were collected at 1 and 24 h after the onset of ^22^Na treatment. After the seedling roots were rinsed with the modified MGRL solution, ^22^Na activity was measured using imaging plates (BAS-IP MS, FUJIFILM, Tokyo Japan) and a Typhoon FLA-7000 image reader (Cytiva, Tokyo, Japan).

### Gene expression analysis

Total RNA extraction and RT-qPCR analysis were generally performed as previously described (Nagatoshi et al. 2023). Cotyledons, hypocotyls, and roots of seedlings treated as described for Na^+^ and K^+^ ion content measurements were rapidly frozen in liquid nitrogen. RNA was extracted from tissues using RNAiso plus (Takara). First-strand cDNA was synthesized from half a volume of total RNA (1 μg) treated for 30 min at 37°C with RQ1 RNase-free DNase (Promega) using a SuperScript III First-Strand Synthesis System (Invitrogen). Gene expression was quantified on a QuantStudio 7 Frex real-time PCR system (Applied Biosystems) using GoTaq qPCR Master Mix (Promega). The primers used in this study are described in Supplementary Table S5.

### RNA-seq analysis

Three quinoa inbred lines (Kd, J100, and J075) grown in vermiculite culture for 10 d were treated with 0 or 600 mM NaCl solution for 0, 6, and 24 h. Total RNA was extracted from cotyledons and roots and treated with DNase as described above. The libraries were prepared and sequenced at Macrogen Japan. The integrity of the total RNA was evaluated using an Agilent 2100 Bioanalyzer (Agilent). Paired-end sequencing (151-bp) of the mRNA library prepared using a TruSeq standard mRNA library Prep was performed on a NovaSeq 6000. All paired-end reads were trimmed using a Trimmomatic (v.0.38) (Bolger et al. 2014) to remove adapter sequence (ILLUMINACLIP:2:20:10) and low-quality reads (SLIDINGWINDOW:4:20, MINLEN:40). The filter-passed reads were mapped on reference sequence from CoGe database (id60716) (Rey et al. 2023) using Hisat2 (v2.1.0) (Kim et al. 2019). Reads mapped to transcript regions were counted and transcripts per million (TPM) values of transcripts were calculated for each sample.

### Differential gene expression analysis

DEGs were identified by organ-specific comparison of cotyledons and roots for 6 and 24 h of salt stress treatment. We performed quantitative analyses using three biological replicates. The statistical difference of read count data between the 0 and 600 mM NaCl treatments in each line was calculated using the edgeR package (Chen et al. 2016). Gene expression was considered significant if the FDR was less than 0.01 and |log_2_(FC)| was greater than 1 in at least one comparison. The resulting gene expression level data obtained were then imported into R software for hierarchical clustering. Prior to the clustering analysis, the TPM values were converted to z-scores for each gene using the “genescale” function included in the “genefilter” package obtained from bioconductor.org. Hierarchical clustering analysis and drawing of heat maps with dendrograms were performed using the heatmap.2 function of the “gplots” package (Warnes et al. 2016).

### Functional enrichment analysis of DEGs in each cluster

GO enrichment analysis was performed with topGO using the classical algorithm with Fisher’s test (Alexa and Rahnenführer 2009). GO annotations were also assigned to the quinoa reference genomes using the Trinotate pipeline (Bryant et al. 2017). The top 50 biological process GO terms significantly enriched in the gene list for each cluster were filtered by applying a 0.05 cutoff to Fisher’s weighted *P*-values.

### Whole-genome resequencing

We used total genomic DNA from 17 quinoa inbred lines for whole-genome resequencing (Mizuno et al. 2020). DNA libraries were prepared using an Illumina TruSeq DNA PCR-Free Kit and sequenced as 151-bp paired-end reads on an Illumina NovaSeq 6000 system at Macrogen Japan. Low-quality reads and adapters were trimmed using Trimmomatic (v.0.38) (Bolger et al. 2014) with the options ‘SLIDINGWINDOW:4:25’ and ‘MINLEN:40’. The trimmed paired-end reads were aligned to the quinoa reference genome (QQ74, v2, id60716) from CoGe (https://genomevolution.org/coge/) using BWA 0.7.17 (Li, 2013). Samtools (v1.8) (Danecek et al. 2021) was used to convert the alignment SAM files to BAM files and prepare the alignment file for viewing in the Integrative Genomics Viewer (IGV) (Robinson et al. 2011).

### Molecular cloning and plasmid construction

Based on the coding sequences (CDSs) for putative *CqHKT1;1, CqHKT1;2* and *CqSOS1* genes obtained from the NCBI database of *Chenopodium quinoa*, which the amino acid sequences of *Arabidopsis thaliana* AtHKT1 (AT4G10310) and AtSOS1 (AT2G01980) were used as queries in the BLASTP program, we designed PCR primers (Supplementary Table S5) to amplify trigger regions for VIGS in the CDSs. ALSV-RNA2 vectors for VIGS analyses were constructed as previously described (Ogata et al. 2017), with minor modifications. The amplified DNA fragments containing trigger sequences of quinoa genes for VIGS were cloned in-frame into the *Xho*I/*Bam*HI site of pEALSR2 to generate pEALSR2-CqHKT1;1, pEALSR2-CqHKT1;2 and pEALSR2-CqSOS1, respectively.

### Virus inoculation and salt stress treatment

ALSV inoculations were performed as previously described (Ogata et al. 2021). The plasmids for the ALSV-RNA1 (pEALSR1) and ALSV-RNA2 constructs were mixed in equal amounts, and the DNA solution was mechanically inoculated onto the any true leaves of 14-day-old quinoa plants (Iw inbred line) using carborundum (Li et al. 2004). The inoculated quinoa plants were grown for 2 to 3 weeks and the uninoculated upper leaves showing chlorotic spots symptoms were harvested. The detached leaves were ground using the ShakeMaster and 3-mm stainless steel beads and suspended in extraction buffer (0.1 M Tris-HCl, pH 8.0, 0.1 M NaCl, 5 mM MgCl_2_) (Igarashi et al. 2009). Debris was precipitated by centrifugation at 18,800 *g* for 10 min at 4°C, and the supernatants were used to inoculate quinoa Kd lines grown in the temperature-controlled phytotron as described above. Infected leaves or the inocula were stored at −80°C prior to use. ALSV derived from quinoa Iw plants inoculated with a mixture of pEALSR1 and pEALSR2 was used as a control (ALSV-WT). Seedlings (14 day old) of quinoa Kd lines were mechanically inoculated with the inocula using carborundum. Kd plants at 10 days post inoculation were treated with a 0.5 L of 300 mM NaCl solution once a week for 3 weeks. For Na^+^ content measurements, the lower leaves above the ALSV-inoculated leaves were collected and oven-dried at 60°C for 2 days. In the same plants, other leaves at similar positions were collected for gene expression analysis. Na^+^ content measurement and gene expression analysis were performed as described above.

### Statistical analysis

All data are presented as mean ± standard deviation. Significant differences in data for physiological and morphological indices were determined by one-way analysis of variance and Tukey’s honestly significant difference (HSD) test or Student’s *t*-test using R version 4.3.0. The data were visualized using the “ggplot2” packages in R.

### Accession numbers

The Illumina read sequencing data from the RNA-seq and whole-genome resequencing were deposited in the DDBJ Sequence Read Archive under Bioproject accession numbers PRJDB18326 and PRJDB18355, respectively. Each accession number of the whole-genome resequencing data for quinoa inbred lines is listed in Table S1.

**--**

## Acknowledgments

We thank Prof. Emer. N. Yoshikawa (Iwate Univ.) for kindly providing the ALSV-VIGS vectors and quinoa seeds that are the source of the inbred line Iw. We thank the staff of JIRCAS, M. Toyoshima, N. Hisatomi, Y. Masamura, K. Ozawa, Y. Shirai, N. Saito, Y. Nakamura, Y. Nonoue, Y. Takiguchi, A. Aoyama, Y. Ishino, N. Seko, T. Nada, M. Nozawa, A. Karasawa, and W. Kawakami for their excellent technical assistance. We also thank K. Katsura, Y. Tokura, G. Almanza, R. Oros, I. Molares, A. Bonifacio, W. Rojas, J. Quezada, M. Patricia, Y. Flores for their cooperation in quinoa research. Computations were performed in part on the NIG supercomputer at ROIS National Institute of Genetics.

## Author contributions

Y.K. and Y.F. conceived and designed the research; Y.K. analyzed the data; R.S. performed the radio isotope analysis; Y.K. M.F. and Y.Y. were involved in the genome analysis; Y.K. performed the HPLC analysis with the help of Y.M.; Y.K. conducted the VIGS analysis with the help of T.O.; Y.K. conducted the gene expression analysis with the help of Y.N.; Y.N. and Y.F. are involved in project administration; Y.F. supervised the research; Y.K and Y.F. wrote and revised the manuscript; All authors contributed to the article and approved the submitted version.

## Supplementary data

**Supplementary Fig. S1.** Growth of seedlings of representative quinoa inbred lines in response to high salt stress over time. Ten-day-old seedlings of quinoa inbred lines were treated with 0 or 600 mM NaCl for 24 h. L, lowland lines; NH, northern highland lines; SH, southern highland lines. To facilitate understanding of the relationship with Na^+^ accumulation, the lines within each genotype are arranged from left to right in order of Na^+^ content in the cotyledons, as shown in Supplementary Fig. S2. Data show mean ± SD of cotyledon, hypocotyl, and root dry weights for each line (*n* = 5).

**Supplementary Fig. S2.** Na^+^ accumulation in seedlings of representative quinoa inbred lines in response to high salt stress over time. Ten-day-old seedlings of quinoa inbred lines were treated with 0 or 600 mM NaCl for 24 h. L, lowland lines; NH, northern highland lines; SH, southern highland lines. To facilitate understanding of the relationship with Na^+^ accumulation, the lines within each genotype are arranged from left to right in order of Na^+^ content in the cotyledons, as shown in Supplementary Fig. S2. Data show mean ± SD of Na^+^ content in the cotyledon, hypocotyl, and root of each line (*n* = 5).

**Supplementary Fig. S3.** K^+^ accumulation in seedlings of representative quinoa inbred lines in response to high salt stress over time. Ten-day-old seedlings of quinoa inbred lines were treated with 0 or 600 mM NaCl for 24 h. L, lowland lines; NH, northern highland lines; SH, southern highland lines. To facilitate understanding of the relationship with Na^+^ accumulation, the lines within each genotype are arranged from left to right in order of Na^+^ content in the cotyledons, as shown in Supplementary Fig. S2. Data show mean ± SD of K^+^ content in the cotyledon, hypocotyl, and root of each line (*n* = 5).

**Supplementary Fig. S4.** High salt stress induces the expression of genes related to cell wall organization in the aerial parts of quinoa seedlings of the lowland line, which accumulates Na^+^ in the aerial parts. GO term enrichment analysis of cluster A genes (Figure 7A), which are induced to a greater extent under high salt treatment in the cotyledons of young Kd seedlings than in those of J075 and J100. Each enriched biological process GO term is shown in the directed acyclic graphical model. The box indicates the 10 most enriched terms out of the 50 GO terms shown in Supplementary Table S2. Different colors represent different degrees of enrichment significance, with darker colors indicating higher significance.

**Supplementary Fig. S5.** High salt stress suppresses the expression of genes associated with hypoxia in the aerial parts of quinoa seedlings of the lowland line, which accumulates Na^+^ in the aerial parts. GO term enrichment analysis of cluster B genes (Figure 7A), which include those suppressed to a greater extent in the cotyledons of young Kd seedlings treated with high salt than in those of J075 and J100. Each enriched biological process GO term is shown in the directed acyclic graphical model. The box indicates the 10 most enriched terms out of the 50 GO terms shown in Supplementary Table S2. Different colors represent different degrees of enrichment significance, with darker colors indicating higher significance.

**Supplementary Fig. S6.** High salt stress suppresses the expression of photosynthesis-related genes in the aerial parts of quinoa seedlings from the lowland line, which accumulates Na^+^ in the aerial parts. GO term enrichment analysis of cluster C genes (Figure 7A), which include those suppressed to a greater extent in the cotyledons of young Kd seedlings treated with high salt than in those of J075 and J100. Each enriched biological process GO term is shown in the directed acyclic graphical model. The box indicates the 10 most enriched terms out of the 50 GO terms shown in Supplementary Table S2. Different colors represent different degrees of enrichment significance, with darker colors indicating higher significance.

**Supplementary Fig. S7.** High salt stress induces the expression of genes associated with responses to water deprivation and ABA in the roots of quinoa seedlings from all three genotypic lines. GO term enrichment analysis of cluster D genes (Figure 7B) shows that all three genotypic lines contain more induced genes in the roots of young seedlings treated with high salt than in untreated roots. Each enriched biological process GO term is shown in the directed acyclic graphical model. The box indicates the 10 most enriched terms out of the 50 GO terms shown in Supplementary Table S2. Different colors represent different degrees of enrichment significance, with darker colors indicating higher significance.

**Supplementary Fig. S8.** High salt stress greatly induces the expression of genes associated with responses to hypoxia in the roots of quinoa seedlings of southern highland line, which does not accumulate much Na^+^ in the aerial parts. GO term enrichment analysis of cluster E genes (Figure 7B), which are induced by high salt treatment to a greater extent in the roots of young J100 seedlings than in those of J075 and Kd. Each enriched biological process GO term is shown in the directed acyclic graphical model. The box indicates the 10 most enriched terms out of the 50 GO terms shown in Supplementary Table S2. Different colors represent different degrees of enrichment significance, with darker colors indicating higher significance.

**Supplementary Fig. S9.** High salt stress induces the expression of genes associated with the defense response to bacteria and hydrogen peroxide catabolic process to a greater extent in the roots of quinoa seedlings of highland lines than in those of a lowland line. GO term enrichment analysis of cluster F genes (Figure 7B), which are induced to a greater extent in the roots of young seedlings treated with high salt in highland lines, J075 and J100, than in those of the lowland line Kd. Each enriched biological process GO term is shown in the directed acyclic graphical model. The box indicates the 10 most enriched terms out of the 50 GO terms shown in Supplementary Table S2. Different colors represent different degrees of enrichment significance, with darker colors indicating higher significance.

**Supplementary Fig. S10.** Transcriptome profiling of quinoa HKT transporter and Na^+^/H^+^ exchanger genes in cotyledons and roots of representative quinoa inbred lines. Heatmap showing hierarchical clustering of TPM of HKT transporter and Na^+^/H^+^ exchanger genes in the cotyledons and roots of seedlings subjected to control conditions (0 mM NaCl) and NaCl treatment (600 mM NaCl) for 0, 6, and 24 h.

**Supplementary Fig. S11.** Insertion of a single adenine in the first exon in the 5′ region of the *CqHKT1;1* gene of the northern highland lines J072, J073, and J075 shifts the reading frame. (A) Visualization of a polymorphism pattern in the *CqHKT1;1* gene in northern highland lines. Read coverage (upper) and alignment (lower) for each genotype were viewed using IGV. Colored vertical lines in the reads represent single nucleotide polymorphisms (SNPs) and insertion-deletion polymorphisms (InDels) detected in the WGS data. Gene models of *CqHKT1;1* are shown in blue below the reads. Reads and their physical locations were aligned to the QQ74 reference genome (v2, id60716). (B) Two sequences at the top and bottom are genomic DNA and amino acid sequences based on the standard rules of genetic decoding of the reference line and northern highland lines (J072, J073, and J075). The insertion at position +68 in an oligoA stretch of *CqHKT1;1* in J072, J073 and J075 is marked in red. Blue letters indicate the start codon.

**Supplementary Fig. S12.** Relative expression levels of transporter genes in response to high salt stress. Ten-day-old seedlings of quinoa inbred lines were treated with 0 or 600 mM NaCl for 24 h. After the treatments, gene expression in cotyledons was examined for *CqHKT1;2*, which is mainly expressed in cotyledons, and gene expression in roots was examined for *CqHKT1;1* and *CqSOS1*, which are mainly expressed in roots. L, lowland lines; NH, northern highland lines; SH, southern highland lines. To facilitate understanding of the relationship with Na^+^ accumulation, the lines within each genotype are arranged from left to right in order of Na^+^ content in the cotyledons, as shown in Supplementary Fig. S2. The transcript levels of *CqHKT1;2*, *CqHKT1;1*, and *CqSOS1* genes were normalized to those of *CqUBQ10* as an internal control gene. Relative expression levels of *CqHKT1;1*, *CqHKT1;2*, and *CqSOS1* are shown as dots relative to the gene expression levels in each corresponding tissue in the non-salt-treated Kd seedlings. Error bars indicate SD (*n* = 5). Relative expression data for *CqHKT1;1* genes in J072, J073, and J075 are presented as “no gene” (*) because an insertion at position +68 in an oligoA of the gene results in a premature stop codon (Supplementary Fig. S11). Relative expression data for *CqSOS1* genes in J064 and J071 are also considered “no gene” (**) due to an approximately 6.9-kb deletion in the 5′ region of the gene (Supplementary Fig. S13).

**Supplementary Fig. S13.** Deletion of the 5′ portion of the *CqSOS1* gene in chromosome 6A of the northern highland inbred lines J064 and J071 disrupts the gene. Visualization of the polymorphism pattern in the *CqSOS1* gene in northern highland inbred lines. Read coverage (upper) and alignment (lower) for each genotype were viewed using IGV. Colored vertical lines in the reads represent SNPs and InDels detected in the WGS data. Gene models of *CqSOS1* are represented in blue below the reads. Reads and their physical locations were aligned to the QQ74 reference genome (v2, id60716).

**Supplementary Fig. S14.** Visualization of the polymorphism pattern in the upstream region of *CqHKT1* genes on chromosome 7B in selected northern highland (NH), southern highland (SH) and lowland (L) inbred lines. Read coverage (upper) and alignment (lower) for each genotype were viewed using IGV. Colored vertical lines in the reads represent SNPs and InDels detected in the WGS data. Gene models of *CqHKT1;1* and *CqHKT1;2* are shown in blue below the reads. Reads and their physical locations were aligned to the QQ74 reference genome (v2, id60716).

**Supplementary Fig. S15.** Visualization of the polymorphism pattern in the upstream region of *CqSOS1* genes on chromosome 6A in selected northern highland (NH), southern highland (SH) and lowland (L) inbred lines. Read coverage (upper) and alignment (lower) for each genotype were viewed using IGV. Colored vertical lines in the reads represent SNPs and InDels detected in the WGS data. Gene models of *CqSOS1s* are shown in blue below the reads. Reads and their physical locations were aligned to the QQ74 reference genome (v2, id60716).

**Supplementary Fig. S16.** Visualization of the polymorphism pattern in the upstream region of *CqSOS1* genes on chromosome 6B in selected northern highland (NH), southern highland (SH) and lowland (L) inbred lines. Read coverage (upper) and alignment (lower) for each genotype were viewed using IGV. Colored vertical lines in the reads represent SNPs and InDels detected in the WGS data. Gene models of *CqSOS1s* are shown in blue below the reads. Reads and their physical locations were aligned to the QQ74 reference genome (v2, id60716).

**Supplementary Table S1.** List of quinoa inbred lines used in this study.

**Supplementary Table S2.** Number of DEGs in representative quinoa inbred lines.

**Supplementary Table S3.** Biological process Gene Ontology terms identified in cotyledons and enriched in each cluster shown in Figure 7A.

**Supplementary Table S4.** Biological process Gene Ontology terms identified in roots and enriched in each cluster shown in Figure 7B.

**Supplemental Table S5.** Primers used in this study.

## Data availability

The data generated in this study are available in the article and in its online supplementary materials.

